# Functional benefits of continuous vs. categorical listening strategies on the neural encoding and perception of noise-degraded speech

**DOI:** 10.1101/2024.05.15.594387

**Authors:** Rose Rizzi, Gavin M. Bidelman

**Author notes:** **Author contributions:**. R. R. and G.M.B. designed the experiment, R.R. collected the data, R.R. and G.M.B. analyzed the data and wrote the paper. **Address for editorial correspondence:** Gavin M. Bidelman, Ph.D. Speech, Language and Hearing Sciences 2631 East Discovery Parkway Bloomington, IN 47408.

## Abstract

Acoustic information in speech changes continuously, yet listeners form discrete perceptual categories to ease the demands of perception. Being a more continuous/gradient as opposed to a discrete/categorical listener may be further advantageous for understanding speech in noise by increasing perceptual flexibility and resolving ambiguity. The degree to which a listener’s responses to a continuum of speech sounds are categorical versus continuous can be quantified using visual analog scaling (VAS) during speech labeling tasks. Here, we recorded event-related brain potentials (ERPs) to vowels along an acoustic-phonetic continuum (/u/ to /a/) while listeners categorized phonemes in both clean and noise conditions. Behavior was assessed using standard two alternative forced choice (2AFC) and VAS paradigms to evaluate categorization under task structures that promote discrete (2AFC) vs. continuous (VAS) hearing, respectively. Behaviorally, identification curves were steeper under 2AFC vs. VAS categorization but were relatively immune to noise, suggesting robust access to abstract, phonetic categories even under signal degradation. Behavioral slopes were positively correlated with listeners’ QuickSIN scores, suggesting a behavioral advantage for speech in noise comprehension conferred by gradient listening strategy. At the neural level, electrode level data revealed P2 peak amplitudes of the ERPs were modulated by task and noise; responses were larger under VAS vs. 2AFC categorization and showed larger noise-related delay in latency in the VAS vs. 2AFC condition. More gradient responders also had smaller shifts in ERP latency with noise, suggesting their neural encoding of speech was more resilient to noise degradation. Interestingly, source-resolved ERPs showed that more gradient listening was also correlated with stronger neural responses in left superior temporal gyrus. Our results demonstrate that listening strategy (i.e., being a discrete vs. continuous listener) modulates the categorical organization of speech and behavioral success, with continuous/gradient listening being more advantageous to speech in noise perception.

## 1. Introduction

Listeners are often tasked with understanding speech signals in noisy listening environments. Speech-in-noise (SIN) perception is a difficult cognitive process and a common audiologic complaint. While certain clinical populations, such as those with hearing loss (Picard et al., 1999; Plomp, 1978), cognitive deficits (Bradlow et al., 2003; Grady et al., 1989), traumatic brain injury (Hoover et al., 2017; Vander Werff & Rieger, 2019), and old age (Bergman, 1971; Humes et al., 2013), show exacerbated SIN difficulty, even normal hearing listeners can have deficits in SIN comprehension (Bharadwaj et al., 2015; Hannula et al., 2011; Ruggles et al., 2011; Tremblay et al., 2015). This large variability in SIN perception emphasizes the importance of analyzing individual differences in performance to understand how a listener’s perceptual strategy might influence SIN outcomes.

Listeners have simultaneous access to both acoustic (continuous) and phonetic (categorical) cues of speech (Andruski et al., 1994; Blumstein et al., 2005; McMurray et al., 2002; Miller & Volaitis, 1989; Pisoni & Tash, 1974). Different listeners may weigh information from these modes differently during speech perception tasks, such that some listeners are categorical/discrete responders, while others are more continuous/gradient responders (Kapnoula et al., 2021; Kapnoula et al., 2017; Kong & Edwards, 2016). Theoretically, either strategy could benefit perception. First, categorical/discrete listening may be an ideal strategy for SIN perception. While acoustic information changes continuously, listeners bin speech sounds into equivalency categories to map speech acoustics to a high-level phonetic code (Liberman et al., 1967; Pisoni, 1973). This more abstract categorical code might be more resistant to degradation by noise, since it does not rely on surface features of speech that are easily washed out by noise (Bidelman et al., 2020; Bidelman et al., 2019). Indeed, more discrete listeners show less interference from informational masking in auditory streaming tasks that mimic naturalistic “cocktail party” listening scenarios (Bidelman et al., 2024). Previous work using event-related brain potentials (ERPs) has also demonstrated enhanced P2 responses to noise-degraded speech sounds with clear phonetic identities compared to ambiguous speech sounds that do not carry a clear phonetic label, suggesting the brain does not linearly code changes in acoustics but represents a categorical code (Bidelman et al., 2020; Bidelman et al., 2013; Bidelman & Walker, 2019; Bidelman & Walker, 2017). The categorical organization for speech is thought to involve a network including auditory cortex (Bidelman & Lee, 2015; Bidelman & Walker, 2019; Chang et al., 2010) and higher-order linguistic centers in the inferior frontal gyrus (IFG) (Alho et al., 2016; Myers et al., 2009) that are differentially engaged depending on task difficulty (Carter & Bidelman, 2021) and a listener’s experience (e.g. language or music background; Bidelman & Lee, 2015; Bidelman & Walker, 2019).

Conversely, a gradient/continuous listening strategy might confer a perceptual advantage for SIN processing. Maintaining within-category acoustic information may allow listeners to “hedge” their bets on the speech sounds they hear and resolve ambiguity (Kapnoula et al., 2017). For instance, when speech sound identities are more uncertain having variable voice onset times (VOTs), listeners respond in a more graded fashion (Clayards et al., 2008). Kapnoula et al. (2021) found that gradient listeners were better able to recover from lexical ambiguity in garden path sentences, though SIN perception was not improved. Gradient listeners might also have more flexibility in cue-weighting when making phonetic decisions about acoustic inputs (Kapnoula et al., 2017; Massaro & Cohen, 1983b; Toscano & McMurray, 2010). Neural evidence of gradient processing from ERPs has demonstrated linear changes in the N1 wave (∼100 ms) with changes in VOT (Toscano et al., 2010), earlier in the time course of the response than categorical information seems to be coded (Bidelman et al., 2013). Though, graded speech representations can be maintained in the neural signal for up to 900 ms (Sarrett et al., 2020), even after category abstraction (Bidelman et al., 2013). These findings suggest the brain likely represents and maintains both within- and across-category contrastive information (Toscano et al., 2018). Imaging studies suggest that graded activation to speech cues occur in left superior temporal gyrus (STG) (Myers et al., 2009). Thus, gradient perception may rely more heavily on sensory representations at the level of auditory cortex, while category labelling might recruit higher order linguistic resources downstream (IFG). While it is clear neural responses scale with acoustic-phonetic features of the speech signal (exogenous properties), it is unclear how they are modulated by endogenous properties of the listeners. Here, we ask whether differences in listener strategy can actively modulate the neural encoding of speech and beneficially transfer to SIN processing.

One barrier to studying listening strategy is the common use of two alternative forced choice (2AFC) paradigms in speech categorization tasks. In 2AFC, listeners hear speech sounds sampled from an equidistant acoustic-phonetic continuum (e.g., /da/ to /ga/). Listeners are asked to press a button to report which of two speech sounds they heard in a forced, binary judgment. The slope of the resulting identification function is often taken as a measure of categoricity in the behavior (Bidelman, 2015; Hallé et al., 2004; Sussman, 1993; Werker & Tees, 1987; Xu et al., 2006). While identification curve slopes assessed under 2AFC can be used as a measure of listener strategy, it remains unclear whether shallower slopes reflect more/less gradiency in perception or simply noisier responses (Apfelbaum et al., 2022; Kapnoula et al., 2017; McMurray et al., 2018). Indeed, disordered populations often have shallower slopes in labeling tasks which is usually interpreted as less categorical hearing (Godfrey et al., 1981; Joanisse et al., 2000; Serniclaes et al., 2001; Serniclaes et al., 2005; Sussman, 1993; Werker & Tees, 1987). SIN deficits are also common in many of these disorders (Cunningham et al., 2001; Dole et al., 2012; Elmahallawi et al., 2021; Lagacé et al., 2010; Warrier et al., 2004; Ziegler et al., 2009), suggesting a shared mechanism between SIN abilities and categorical perception. However, it is plausible that 2AFC responses in these populations are simply less consistent, resulting in shallower identification slopes due to internal perceptual noise rather than difficulty forming categories.

More recent work has demonstrated that using a visual analog scale (VAS) yields measurements of listener strategy that are more independent of internal noise (Kapnoula et al., 2017), suggesting this method is a better for quantifying listener strategy. Whereas the 2AFC task promotes categorical reporting, the VAS task allows for more open-ended responses, thereby reflecting more nuances in listeners’ perception of acoustic cues (Munson et al., 2017). For example, VAS response distributions distinguish categorical from gradient listeners in that discrete listeners make more use of the endpoints of the scale and gradient listeners distribute responses across the entire scale (Kapnoula et al., 2017; Kong & Edwards, 2016; Massaro & Cohen, 1983a).

Current literature has not consistently demonstrated a perceptual or neurophysiologic benefit for either categorical or continuous listening on SIN perception. It is also unclear how individuals’ neural responses may be modulated by task demands alone (i.e., promoting gradient responses in a VAS task and categorical responses in a 2AFC task). In the present study, we measured EEG and behavioral responses during a phoneme labelling task under 2AFC and VAS paradigms to quantify listeners’ categoricity/gradiency in perception. We then assessed correspondences between listening strategy, neural responses, and standardized measures of SIN perception. Our results demonstrate better SIN comprehension scores and stronger neural responses in left STG correspond with more gradient listening, establishing a neural and perceptual link between SIN performance and listening strategy.

## 2. Methods

### 2.1 Participants

Our sample included N=20 English-speaking young adults (19-30 years old, 10 female) with 17.5 ± 2.4 years of education and 8.7 ± 8.0 years of self-reported formal music training. Years of musical training did not correlate with listening strategy measures or SIN performance (all *p*s > 0.05). Participants all had normal hearing (≤25 dB HL; 250-8000 Hz octave frequencies) and were mostly right-handed (73% ± 30%; Edinburgh Handedness Inventory; Oldfield, 1971). Participants provided written informed consent in accordance with a protocol approved by the Institutional Review Board at Indiana University and were paid $10 an hour for their time.

### 2.2 Stimuli and Task

Prior to EEG testing, listeners completed the QuickSIN assessment (Killion et al., 2004) to measure individual SIN comprehension abilities. Sentences were presented binaurally over headphones. The average of scores from two lists of QuickSIN sentences was used to determine a listener’s dB SNR loss, reflecting the SNR threshold required for 50%-word recall.

For the EEG experiment, stimuli consisted of 5 synthetic vowel sounds along a continuum from /u/ to /a/ changing in first formant frequency (F1) (Bidelman et al., 2020; Bidelman et al., 2013). Tokens were sampled from evenly spaced points along a continuum changing F1 linearly from 430 Hz to 730 Hz (**Figure 1A**). Tokens had identical F0 (150 Hz), F2 (1090 Hz), and F3 (2350 Hz). Tokens were 100 ms in duration gated with 10 ms ramps. Stimuli were presented using MATLAB (The MathWorks, Natick; MA, USA) coupled to a TDT RZ6 (Tucker-Davis Technologies, Alachua, FL, USA) signal processor at 75 dB SPL binaurally over shielded insert headphones (ER-2; Etymotic Research).

**Figure 1.**
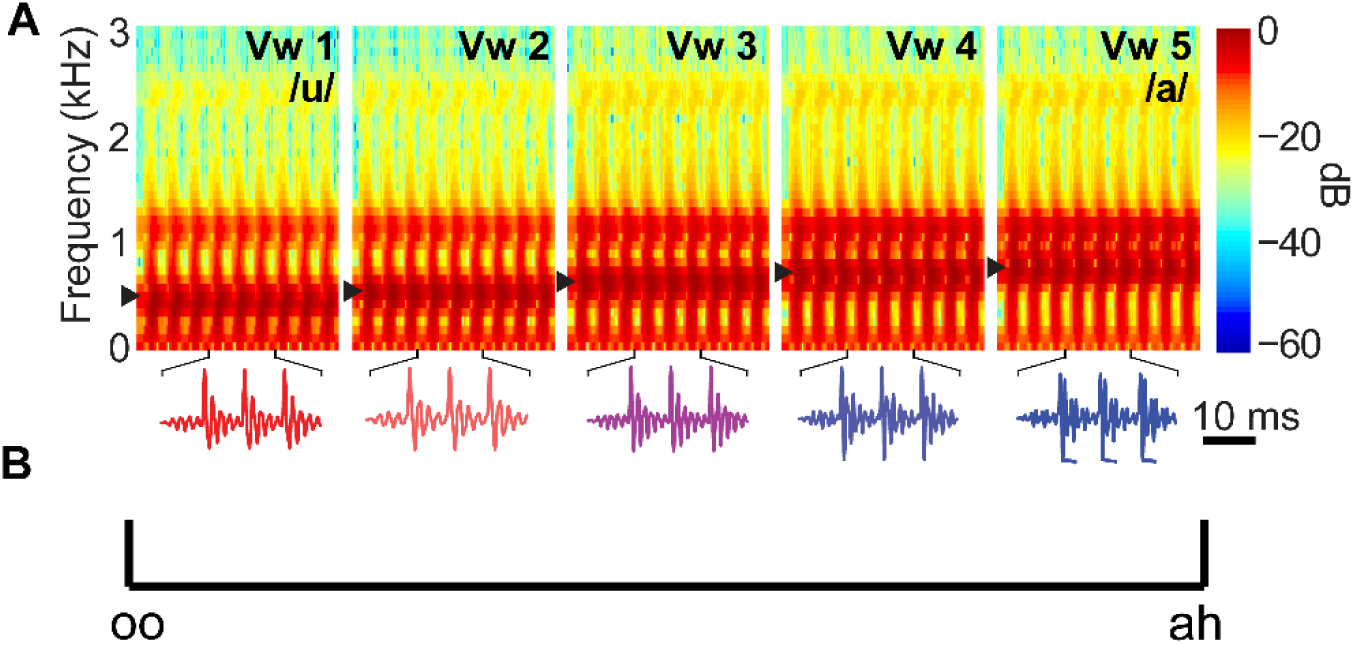
Stimuli and task design. **(A)** Stimulus spectrograms. F1 was changed from 430 to 730 Hz to produce an acoustic-phonetic continuum from “oo” to “ah”. **(B)** Visual analog scale shown to participants during VAS task blocks. Participants were asked to click on the scale to report what they heard.

**Figure 2.**
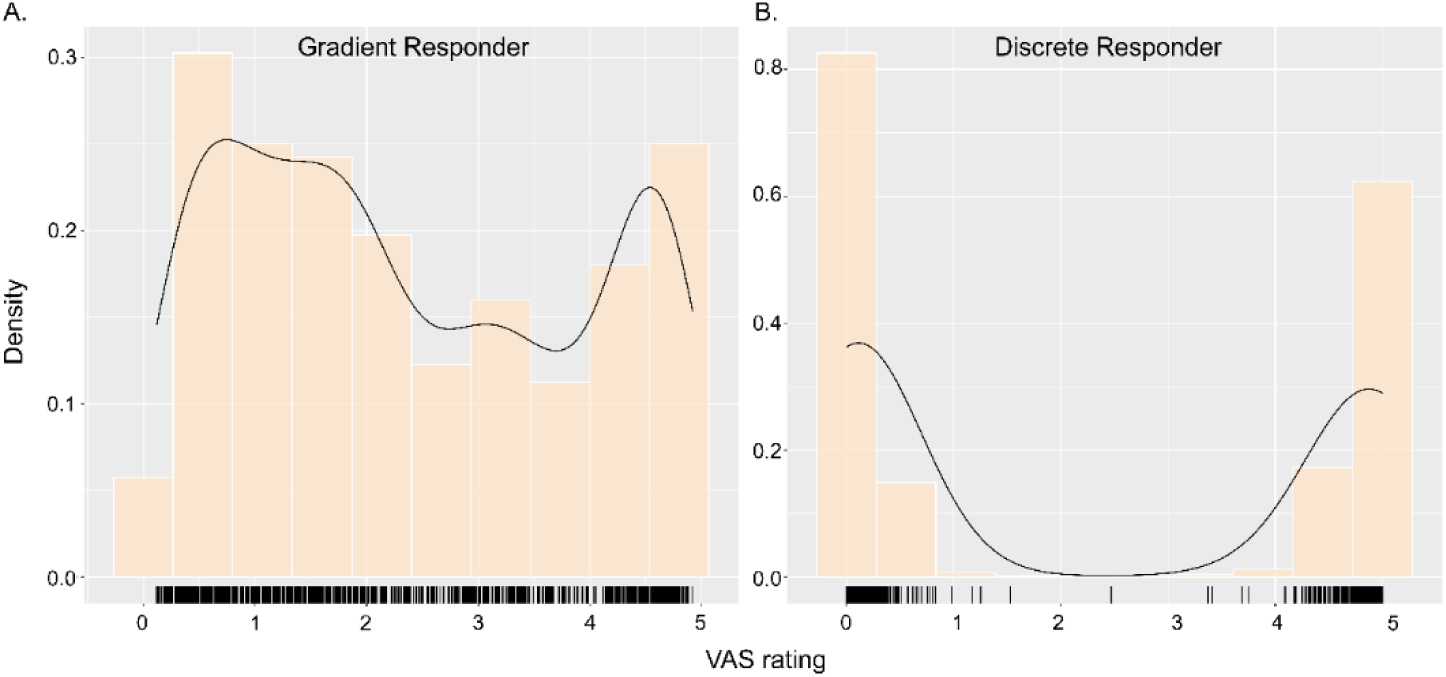
Listeners vary in their response distributions during phoneme labeling. Behavioral response distributions for n=2 representative subjects (clean condition) during VAS labeling. Some listeners report their percept using the entirety of the scale (**A**: gradient responder), while others distribute their responses toward the endpoints of the scale (**B**: discrete responder). Black lines show density plots.

**Figure 3.**
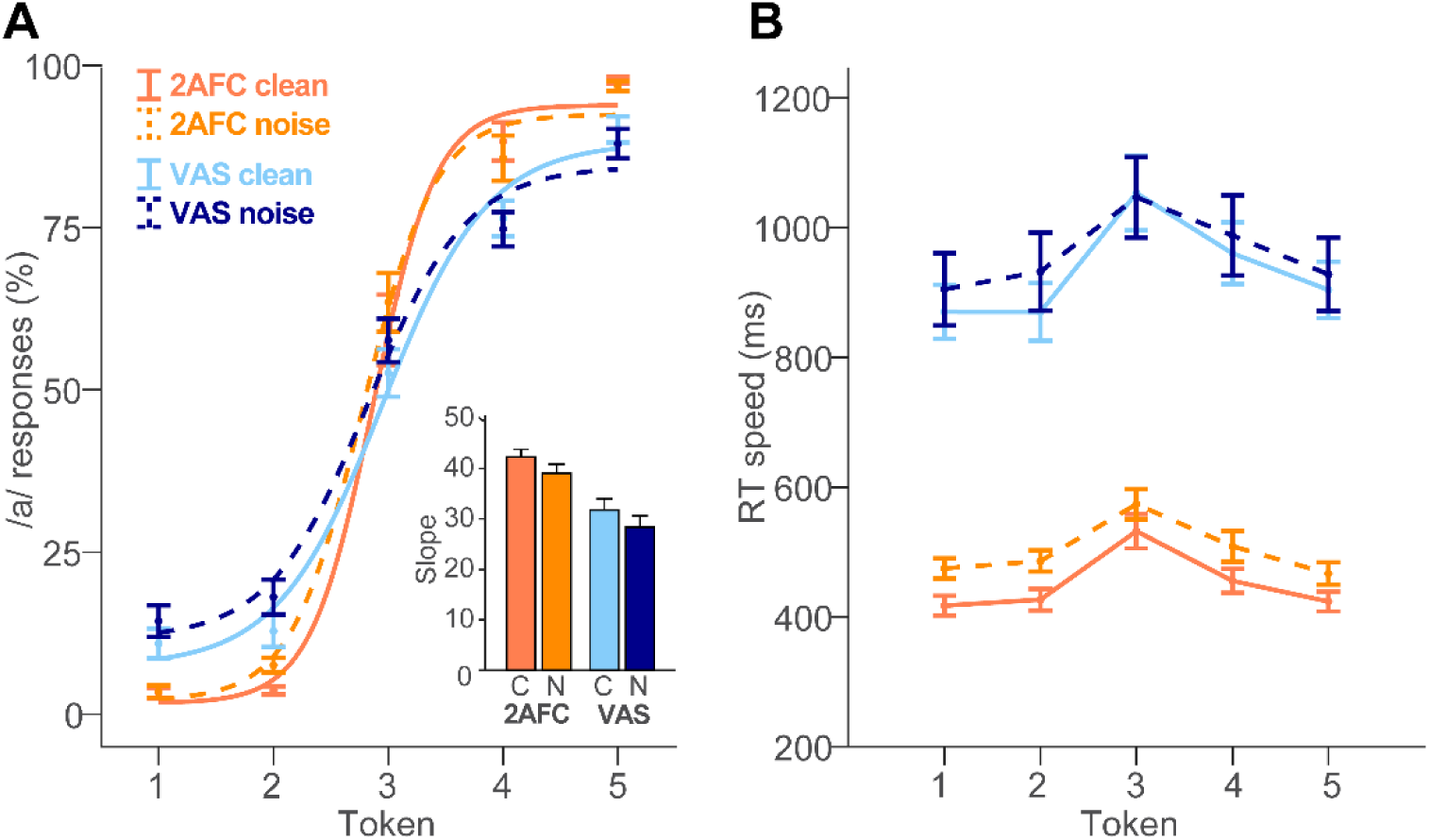
Behavioral identification changes with categorization task and noise conditions. **(A)** Behavioral vowel identification follows stair-stepped function typical for categorical perception. Steeper slopes, indicative of more categorical hearing, were observed in the 2AFC clean condition. Slopes became shallower under VAS labeling and more minimally with the addition of noise. Inset, mean slope in each condition (C=clean, N= noise). **(B)** RTs are slowest for the midpoint (ambiguous) token of the continuum. Faster RTs are observed during 2AFC vs. VAS labeling and (to a lesser extent) for clean relative to noise-degraded speech. Error bars = ± 1 s.e.m.

Vowel stimuli were presented in one of two noise conditions: clean and noise (-2.5 dB SNR). We selected this SNR based on previous findings showing speech categorization is resilient to noise down to ∼0 dB SNR (Bidelman et al., 2020) and pilot testing, that confirmed - 2.5 dB SNR hindered speech perception while still maintaining categorical identification. We used a speech-shaped noise based on the long-term power spectrum (LTPS) of the vowel continuum. Noise was presented continuously throughout the noise block so that it was not time-locked to the stimulus presentation (Alain et al., 2012; Bidelman & Howell, 2016).

During each block, listeners heard 150 presentations of each token and were asked to identify the vowel they heard as quickly and accurately as possible using either a (i) 2 alternative-forced choice (2AFC) binary key press or (ii) visual analog scale (VAS) response. 2AFC and VAS tasks were run in separate blocks but used otherwise identical stimuli; only the task paradigm differed. The VAS paradigm required participants to click a point along a continuous visual scale with endpoints labeled “oo” and “ah” to report their percept (**Figure 1B**). Use of the entire analog scale was encouraged. Following listeners’ behavioral response, the interstimulus interval (ISI) was jittered randomly between 800 and 1000 ms (20 ms steps, uniform distribution) to avoid rhythmic entrainment of the EEG and the anticipation of subsequent stimuli. In total, there were 4 conditions: 2AFC/VAS in clean/noise. Block order was counter-balanced between participants using a Latin square.

### 2.3 Behavioral Data Analysis

To analyze the behavioral responses, we computed the identification curve slopes for each condition, computed as the rise/run change in %-labeling between tokens straddling the midpoint category boundary (i.e., vw2, vw4). Steeper slopes are indicative of more categorical/discrete listening. To provide another quantitative measure of listener strategy, we also used the distribution of VAS responses to calculate Hartigan’s dip statistic, a number that quantifies how multimodal a distribution is (Hartigan & Hartigan, 1985). Higher dip statistic values indicate a more bimodal distribution, representative of a more discrete response strategy (Bidelman et al., 2024). Behavioral speech labeling speeds (i.e., reaction times, RTs) were computed as listeners’ median response latency across trials for a given condition. RTs outside 250–2500 ms were deemed outliers (e.g., fast guesses, lapses of attention) and were excluded from the analysis (Bidelman et al., 2020; Bidelman et al., 2013).

### 2.4 EEG Recording and Data Processing

During each block of behavioral tasks, we recorded high density EEG using 64-channel Ag/AgCl electrodes located at 10-10 positions on the scalp (Oostenveld & Praamstra, 2001). We used Neuroscan Curry 9 software and SynAmps RT Amplifiers (Compumedics Neuroscan, Charlotte, NC) to digitize recordings at 500 Hz. Data preprocessing was then performed in BESA Research 7.1 (BESA, GmbH). During recording, the EEG was referenced to an electrode located 1 cm behind Cz. Recordings were later re-referenced to a common average reference. Single electrodes on the outer canthi of the eyes and the superior and inferior orbit recorded eye movements. We used principal component analysis to correct ocular artifacts (Wallstrom et al., 2004). Additional epochs >150 μV were rejected as artifacts. EEGs were then bandpass filtered from 2 to 30 Hz (zero-phase filters, 48 dB/octave slope). Recordings were then epoched (-200- 800 ms), baselined, and ensemble averaged across trials to generate ERPs for each token per noise and task condition.

### 2.5 ERP Analysis

To reduce the dimensionality of the electrode-level data, ERPs were quantified using 5 electrode clusters (Carter et al., 2022). We averaged activity from adjacent electrodes within each cluster area on the scalp: front left (AF3, F3, F1), front right (AF4, F2, F4), left temporal (FT7, FC5, FC3, T7, C5, C3, TP7, CP5, CP3), right temporal (FC4, FC6, FT8, C4, C6, T8, CP4, CP6, TP8), and center (FC1, FCz, FC2, C1, Cz, C2). ERPs were quantified in latency and amplitude in the time window of the P2 (140-320 ms). We chose to analyze the P2 peak because it occurs in the time course of the ERP when speech categories fully emerge in the brain and it is sensitive to degraded speech perception skills (Bidelman et al., 2020; Bidelman & Lee, 2015; Bidelman et al., 2013; Bidelman & Walker, 2017; Ross et al., 2013). This window was based on visual inspection of the grand average waveform and ensured we captured both noise- and task-related related shifts in P2 latency (Bidelman et al., 2020) (see Figs. 4-5)

**Figure 4.**
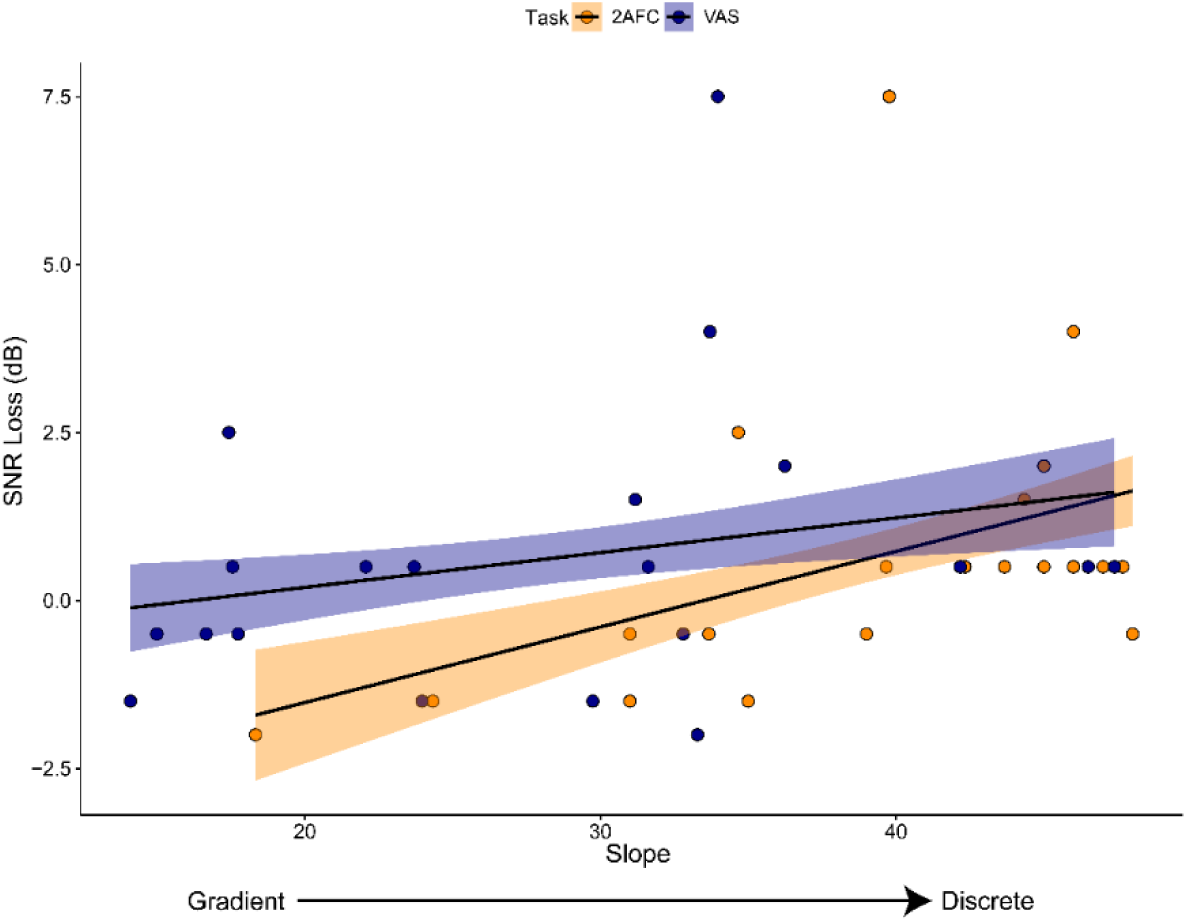
Successful SIN performance (sentence-level recognition) is predicted by a gradient listening strategy (discrete phoneme identification). Pearson correlations between slope in vowel identification and QuickSIN scores (lower score = better SIN comprehension). Only the noise conditions are shown for clarity. Shading = 95% CI.

**Figure 5.**
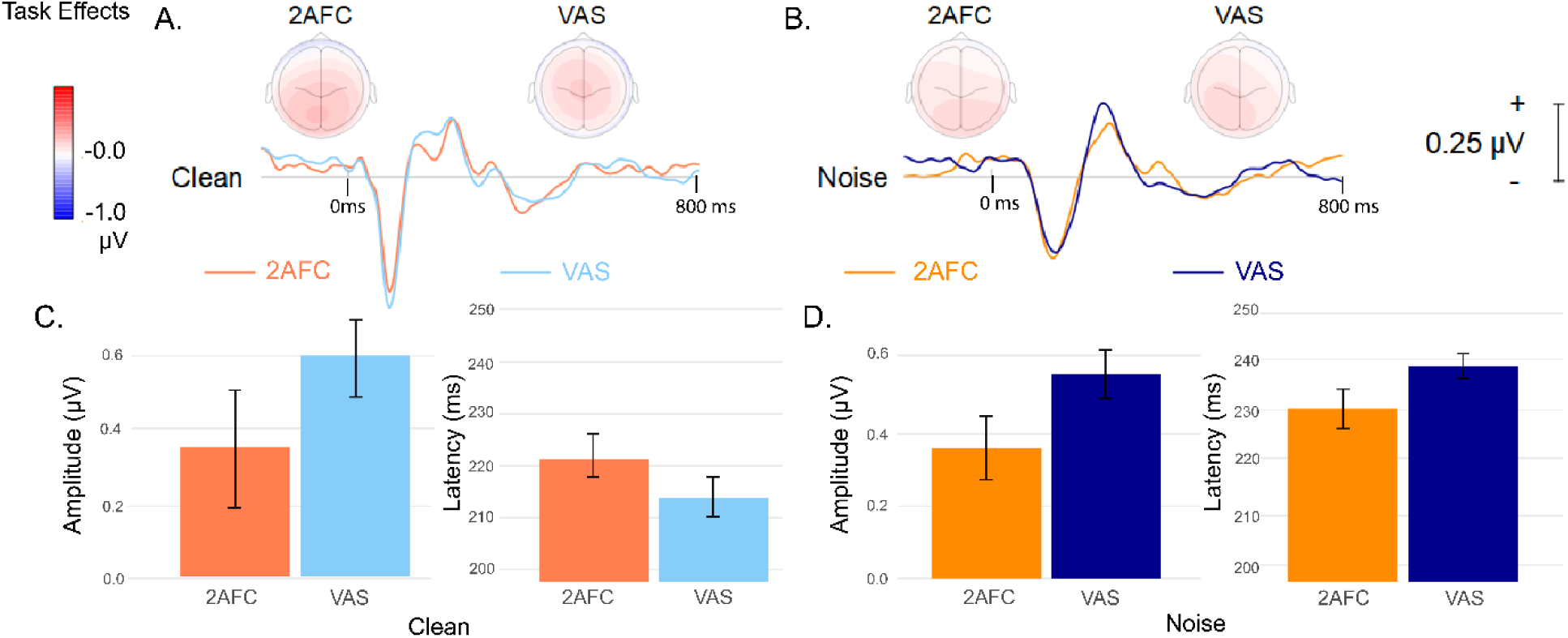
Categorization task modulates cortical responses to otherwise identical speech phonemes. Grand average ERP waveforms measured at the central electrode cluster (FC1, FCz, FC2, C1, Cz, C2) and topographic maps (P2 latency window; 140-320 ms) for **(A)** Clean and **(B)** Noise-degraded speech. **(C-D)** P2 peak amplitudes and latencies (only endpoint tokens [vw1/vw5] shown). Error bars = ± 1 s.e.m.

### 2.6 Statistical Analysis

For behavioral data and ERPs, we used linear mixed model ANOVAs (R; lme4 package; version 1.1-31) to test differences in outcome variables (slope, dip statistic, RT, P2 amplitude, P2 latency). Multiple comparisons were corrected using Tukey-adjusted contrasts with an overall α=0.05. Vowel, task, and noise conditions were fixed effects and subjects served as a random effect. We used Pearson’s correlations to characterize behavior-behavior and brain-behavior relationships. Effect sizes are reported as partial eta squared (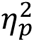) and degrees of freedom were calculated using Satterthwaite’s method.

### 2.7 Source Analysis

We used Classical Low Resolution Electromagnetic Tomography Analysis Recursively Applied (CLARA) [BESA® Research (v7.1)] (Iordanov et al., 2016; Iordanov et al., 2014; Scherg et al., 2019) with a 4-shell ellipsoidal head model (conductivities of 0.33 [brain], 0.33 [scalp], 0.0042 [bone], and 1.00 [cerebrospinal fluid] (Berg & Scherg, 1994) to determine the intracerebral sources that account for continuous vs. discrete listening strategies in speech categorization (e.g., Alain et al., 2023; Bidelman et al., 2018; Carter et al., 2022). Source images were computed for endpoint (vw1/vw5) tokens within the 140-320 ms (∼P2 wave) analysis window, where task and noise effects were maximal in the scalp ERPs (see Figs. 4-5). CLARA models the inverse solution as a large collection of elementary dipoles distributed over nodes on a mesh of the cortical volume. The algorithm estimates the total variance of the scalp data and applies a smoothness constraint to ensure current changes minimally between adjacent brain regions (Michel et al., 2004; Picton et al., 1999). CLARA renders more focal source images by iteratively reducing the source space during repeated estimations. On each iteration (x2), a spatially smoothed LORETA solution (Pascual-Marqui et al., 2002) was recomputed and voxels below a 10% max amplitude threshold were removed. This provided a spatial weighting term for each voxel on the subsequent step. Two iterations were used with a voxel size of 7 mm in Talairach space and regularization (parameter accounting for noise) set to 0.01% singular value decomposition. Source activations were visualized on BESA’s adult brain template (Richards et al., 2016), providing a distributed image describing the P2 activation across the entire brain volume.

We used cluster-based permutation tests (Maris & Oostenveld, 2007) implemented in BESA Statistics (2.1) to examine correlations between the neural source and behavior measures and identify anatomical locations within the full-brain volume that predicted listeners’ degree of categorical hearing (see **Fig. 7A**). For each voxel, a Pearson’s correlation was computed between neural (CLARA source activations) and behavioral (identification slopes) responses. Statistical maps were corrected for multiple comparisons across space by building voxel clusters that control the familywise error rate via a Monte-Carlo resampling technique (Maris & Oostenveld, 2007). We used an alpha level of *α*=0.001 and N=1000 permutations for cluster building. This more stringent alpha level allowed for separation from nearby sources. To better visualize the brain-behavior relations, we then extracted peak CLARA activations and latencies from each significant cluster in the brain volume and regressed these values against listeners’ behavioral identification slopes (see **Fig. 7C, D**).

**Figure 6.**
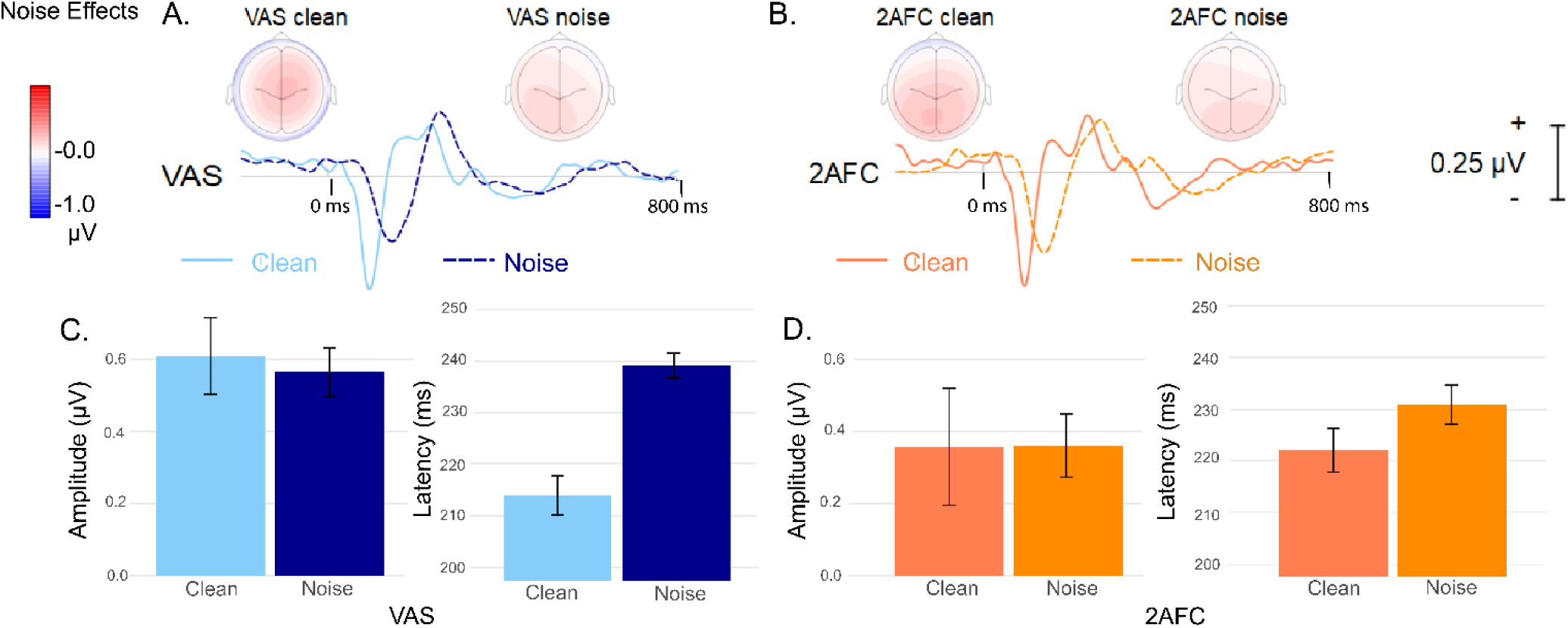
Noise modulates P2 amplitudes and latencies. Note the task x noise interaction depicted by the exaggerated shift in latency in the VAS compared to the 2AFC condition (panel D). Otherwise as in Figure 4. Error bars = ± 1 s.e.m.

**Figure 7.**
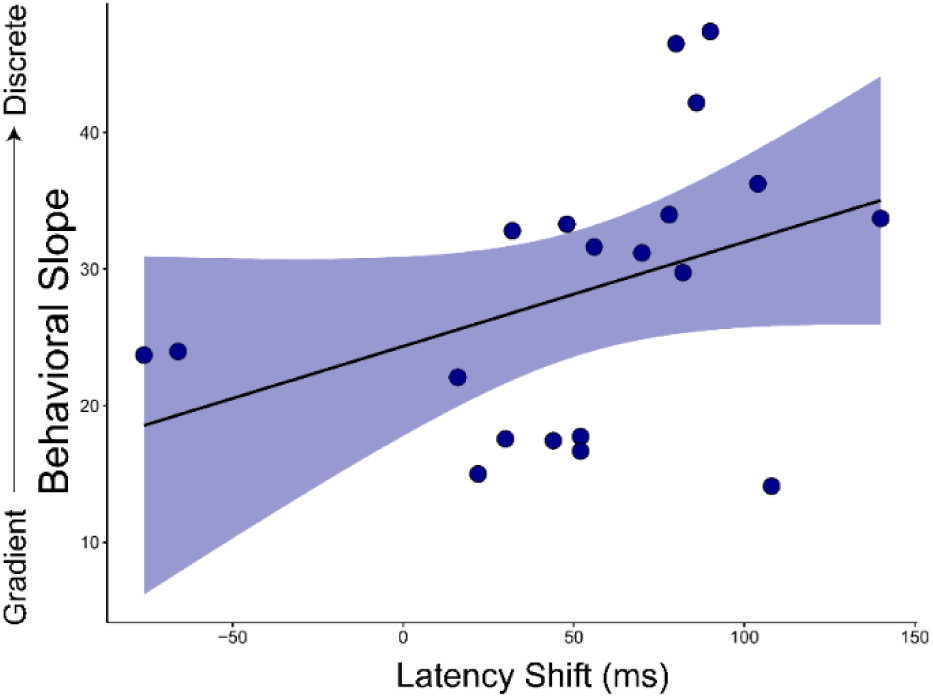
Gradient listeners are more resilient to noise related changes in speech encoding. Latency shift represents the difference in P2 timing elicited in the VAS condition with and without background noise (i.e., P2_noise_ – P2_quiet_). Behavioral slope is positively correlated with noise-related shift in P2 latency. Neural timing is less strongly impacted by noise in more gradient responders. Shading = 95% CI.

## 3. Results

### 3.1 Behavioral Data

We first confirmed listeners’ VAS responses were subject to individual differences and thus showed evidence of gradient vs. discrete listening strategies. **Figure 2** shows responses from n=2 representative subjects who were deemed “gradient/continuous” and “discrete/categorical” responders, respectively. Gradient listeners (**Fig. 2A**) tended to respond along the entire scale to report their percept, while more discrete listeners’ (**Fig. 2B**) responses tended to cluster around the endpoints of the continuum. This confirms listeners do not respond uniformly using the VAS, motivating us to quantify individual differences in their behavioral response patterns.

An ANOVA conducted on identification curve slopes revealed that listeners had steeper slopes (i.e., more categorical labeling) in the 2AFC compared to the VAS task [*F*(1,57) = 49.55, *p* < 0.0001, 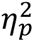 = 0.47] and for clean compared to noise-degraded speech [*F*(1,57) = 4.91, *p* = 0.03, 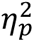 = 0.08] (**Figure 3**). The dip statistic, measured only for the VAS task blocks, decreased with noise [*F*(1,209) = 18.94, *p* < 0.001, 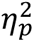 = 0.08]. Behavioral VAS slopes were highly correlated with the dip statistic [*r*(38) = 0.87, *p* < 0.0001], confirming, as expected, more dichotomous responses were associated with bimodal VAS distributions.

Likewise for RT speeds, there was a significant main effect of token [*F*(4,361) = 12.45, *p* < 0.0001, 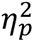 = 0.12], task [*F*(1,361) = 1106.37, *p* < 0.0001, 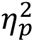 = 0.75], and SNR [*F*(1,361) = 7.91, *p* = 0.005, 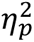 = 0.02]. The task effect was attributed to faster responses under 2AFC vs. VAS labeling whereas the SNR effect was due to slightly faster responses in clean vs. noise. The token effect was attributed to the hallmark slowing in RTs for tokens hear the ambiguous midpoint of the continuum (Bidelman et al., 2020; Bidelman & Walker, 2017; Pisoni & Tash, 1974). This slowing occurred regardless of task or noise SNR [contrast: mean(Tk1,2,4,5) vs. Tk3; all *p*-values < 0.012].

Shallower identification functions could result from weaker categorization and/or noisier responding, both of which would flatten a sigmoid function (Kapnoula et al., 2017). To test the possibility that changes in identification slopes were due to noisier responding, we calculated the standard deviation of subjects’ responses to each token during the VAS conditions, which provides a proxy of sensory noise (Kapnoula et al., 2017). Response noise was not correlated with slope measures [*r*(38) = 0.02, *p* = 0.92], suggesting the steepness of listeners’ identification functions was independent of sensory noise in the decision process (and therefore due to more/less discrete hearing).

Importantly, we found listeners’ QuickSIN scores were highly correlated with categorization slopes for both tasks (2AFC: *r*(38) = 0.36, *p* < 0.0001; VAS: *r*(38) = 0.30, *p* < 0.0001) and SNR conditions (clean: *r*(38) = 0.21, *p* <0.0001; noise: *r*(38) = 0.39, *p* < 0.0001) (**Figure 4**). Listeners with lower dB SNR loss (i.e., better SIN comprehension) had more gradient (shallower) behavioral slopes, suggesting a continuous listening strategy may be more beneficial than a categorical strategy for SIN understanding.

### 3.2 Electrophysiological Data

#### 3.2.1 Electrode Level Data

**Figures 5** and **6** show the electrode-level waveforms, topographic maps, and P2 amplitudes and latencies at the central electrode cluster. P2 amplitudes at the central cluster differed across vowels [*F*(5,437) = 3.19, *p* = 0.0078, 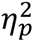 = 0.04], SNR [*F*(1,437) = 54.89, *p* < 0.001, 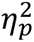 = 0.11], and task [*F*(1,437) = 22.5, *p* < 0.001, 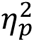 = 0.05]. VAS amplitudes were larger than 2AFC amplitudes [*t*(437) = -4.743, *p* <0.001] across noise conditions. There was also a main effect of noise on P2 latencies [*F*(1,437) = 138.55, *p* < 0.001, 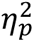 = 0.24]. Interestingly, P2 latencies showed a SNR x task interaction [*F*(1,57) = 6.43, *p* = 0.012, 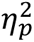 = 0.01], whereby VAS latencies were more prolonged by noise [*t*(437) = -10.12, *p* <0.001] than 2AFC latencies [*t*(437) = -6.53, *p* <0.001]. Similar effects were found at the other electrode clusters (all *p*-values < 0.01) with an additional main effect of vowel at all but the center cluster (all *p*s < 0.0001).

To further investigate the task x SNR interaction, we examined whether noise-related changes in the ERPs predicted the degree of listeners’ behavioral categorization. To this end, we correlated P2 latency shifts produced by noise (i.e., P2_noise_ – P2_quiet_) during the VAS task with listeners’ behavioral slopes. We found that more gradient responders had smaller noise-related P2 latency shifts, suggesting their speech ERPs were more resilient to noise-degradation [*r*(18) = 0.48, *p* = 0.032] (Figure 7)^1^. These findings imply that neural timing to speech was less strongly impacted by noise in more gradient responders.

#### 3.2.2 Source Level Data

To resolve the underlying sources that might contribute to these scalp effects, we examined source-resolved activity using CLARA distributed imaging (Carter et al., 2022; Iordanov et al., 2016). Figure 8A shows correlations between the raw source activations in the P2 time window and behavioral identification across the full brain volume. The cluster analysis and permutation statistics procedure revealed one significant (*p* = 0.012) spatiotemporal cluster encompassing auditory-sensory regions within left superior temporal gyrus (_L_STG; Talairach coordinates: x = -52.5, y = -23.9, z = 2.7) (Fig. 8B). Both the peak amplitude (*r* = -0.46, *p* <0.0001; Fig. 8C) and latency (*r* = 0.26, *p* = 0.021; Fig. 8D) of these _L_STG source activations were strongly correlated with the listeners’ perception. That is, weaker and later STG responses were associated with steeper identification slopes and thus more categorical (discrete) hearing. Conversely, stronger and earlier STG responses were associated with shallower speech labeling and thus more continuous modes of listening.

**Figure 8:**
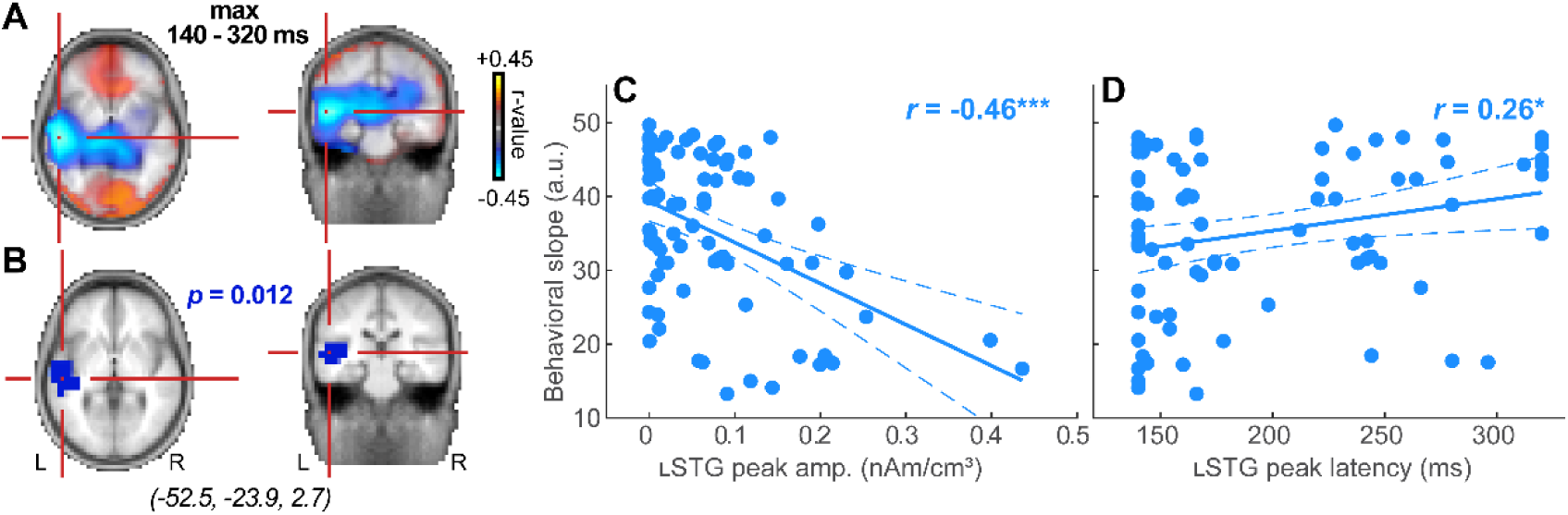
Source localization of neural responses distinguishes discrete vs. continuous listening strategies in speech categorization. (**A**) Full-brain correlations between CLARA source activations and behavioral identification slopes. Hot/cool colors= positive/negative associations. Statistical maps are projected onto BESA’s standard brain template. (**B**) Cluster-based permutation statistics identified a single cluster in left STG driving the brain-behavior relation (cross-hair; Talairach: x = -52.5, y = -23.9, z = 2.7). Both peak amplitude (**C**) and latency (**D**) of STG source activity predict the degree of listeners’ categorical hearing. Weaker/later responses are associated with more discrete phonetic labeling, and stronger/earlier responses with more gradient hearing. Dotted lines = 95% CI, \**p*<0.05, \*\*\**p*<0.0001.

## 4. Discussion

Prior work has been equivocal on whether a categorical or continuous listening strategy is optimal for degraded speech perception. By measuring ERPs to clean and noise-degraded vowel sounds during behavioral tasks that require more/less continuous vs. categorical hearing, we investigated the neural mechanisms subserving the relationship between listening strategy and SIN performance. Subjects’ identification curve slopes served as a measure of listening strategy independent of internal noise. We found continuous listeners, with shallower behavioral slopes and more gradient phoneme perception, performed better on the QuickSIN, suggesting a perceptual advantage for SIN comprehension afforded by a continuous listening strategy. Our data also reveal that speech-ERPs are modulated by task structure, such that otherwise identical speech sounds are differentially encoded by the brain depending on post-perceptual task demands. We also found prominent noise-related changes in ERPs that depended on listening strategy; gradient listeners were more resilient to noise than their discrete listening peers. Lastly, source analysis revealed these neural effects were attributed to changes in _L_STG activation, whereby gradient listening was associated with larger and faster responses in this early, left lateralized auditory region. Collectively, our results demonstrate both behavioral and neural benefits of a gradient listening strategy to speech sound processing and SIN perception.

### 4.1 Speech categories are robust to noise

Behaviorally, we found categorization remained strong in noise with only a small reduction in behavioral slopes at more difficult SNRs. This supports behavioral findings from Bidelman et al. (2020) showing categorical representations for speech persist even in perceptually challenging levels of noise. These results suggest that binning speech sounds into categories is a robust perceptual process and may confer advantages to SIN perception (Bidelman et al., 2020; Bidelman & Carter, 2021). Higher-level category representations can be maintained even when acoustic representations are degraded by noise. RTs followed classic slowing in labelling around the ambiguous midpoint of the continuum in both task and noise conditions; responses were fastest for speech sounds carrying a clear phonetic label (Bidelman & Walker, 2017; Pisoni & Tash, 1974). This suggests, in the broadest sense, category speech presentations might provide an easy, faster readout to perceptual processing. RT delays in the VAS relative to 2AFC task are attributable to differences in the behavioral response requiring the use of a mouse click rather than a button press (Bidelman et al., 2024). Still, the question under investigation here was how these categorization skills actually relate to complex SIN processing.

### 4.2 Categorization skills are related to SIN perception but show an advantage for gradient listening strategies

Notably, we found that more gradient listeners (as measured via phonetic categorization), achieved better SIN comprehension scores (measured via QuickSIN). Gradient listeners may be better equipped to deal with uncertainty by maintaining within-category information to better “hedge” their bets when a speech signal is ambiguous (Kapnoula et al., 2017; McMurray et al., 2008), such as during SIN testing.

Prior work has failed to establish a consistent correlation between SIN performance and listening strategy. This discrepancy is likely due to differences in experimental task design. Neural and behavioral evidence from phoneme labelling tasks in Bidelman et al. (2020) suggests that stronger categorization (i.e., more discrete listening strategy) could aid in successful SIN performance. However, SIN measures in that study were based solely on a phoneme labelling in noise task (2AFC), whereas the present study used more complex SIN assays via the QuickSIN. Moreover, our use of VAS labeling to evaluate listener strategy with regard to categorization skills provides a cleaner measure than 2AFC as it is less obstructed by internal noise that can confound the interpretation of identification curve slope data (Kapnoula et al., 2017). Collectively, our data support the notion that gradient/continuous listening strategies are more beneficial to real-world SIN listening scenarios.

Our findings differ from several prior studies assessing putative relations between categorization and SIN skills. Kapnoula et al. (2021) sought to correlate success of word comprehension in noise with listening strategy measured by a VAS during a garden path sentences task. Though garden path sentences require listeners to resolve ambiguity, a theoretically similar cognitive process to listening to speech in noise, this task does not directly use degraded speech stimuli. Our study thus differs from Kapnoula et al (2021) in that we directly relate two measures of SIN perception at the phoneme and sentence levels. More akin to the present study, Bidelman et al. (2024) demonstrated that discrete listeners assessed by a VAS task had less interference from informational masking than gradient listeners in cocktail party streaming tasks. While our results conflict with those of Bidelman et al. (2024), it is highly likely that different strategies could be deployed across different listening environments to optimize perceptual performance—a single listening “mode” might not be a one size fits all. While gradient listeners may perform better on sentence recognition tasks in multi-talker babble, auditory streaming tasks may engage separate processes that benefit from a discrete listening strategy (Bidelman et al., 2024). Whether listeners change their perceptual strategies or remain consistent across acoustic environments is not clear. An ideal listener may be one that weighs continuous and categorical cues optimally or even dynamically shifts their weights according to the present listening demands. Future studies using a large variety of ecological SIN assessments are needed to test this possibility.

We used behavioral slopes to index listener categorization strategy. Controversy exists regarding whether the slope measure of a sigmoidal identification function truly reflects listener strategy or rather represents internal noise due to perceptual uncertainty (Kapnoula et al., 2017; McMurray, 2022). That is, a shallower identification function could theoretically index either a more gradient listener or a noisier responder. Kapnoula et al. (2017) and Bidelman et al. (2024) both demonstrated that slopes calculated from VAS responses did not reflect response noise. Similarly, we found that estimates of perceptual noise via response variance were not correlated with behavioral slopes. This suggests that slopes during VAS labeling are a veridical measure of listeners’ categorization strategy independent of sensory noise. If categorization slopes were confounded by internal noise, less noisy responders (i.e., those with steeper slopes) should theoretically be more successful in SIN perception. On the contrary, we actually find the opposite pattern: a positive correlation between QuickSIN scores and behavioral slopes. This reinforces the notion that continuous/gradient categorizers, not noisier responders, *per se*, are more skilled SIN listeners.

### 4.3 Speech-ERPs are modulated by task and noise

Our electrophysiological data showed that P2 evoked by otherwise identical speech stimuli was modulated depending on whether listeners were performing 2AFC or VAS labeling. This suggests that the neural encoding and early sensory representations for speech are altered by task demands. Interestingly, we observed larger responses in the VAS condition compared to the 2AFC condition. To our knowledge, this is the first study to demonstrate changes in the auditory-sensory ERPs with changes in post-perceptual task structure.

Listeners have access to both categorical and continuous cues simultaneously which can be used differently with varying task demands, evidenced by reaction time (Pisoni & Tash, 1974), eye tracking (Clayards et al., 2008; McMurray et al., 2018), and MRI studies (Fuhrmeister & Myers, 2021). Some tasks, including the 2AFC identification task used here, may rely only on a categorical or phonetic listening mode, while other tasks may require additional access to a more continuous or acoustic listening mode. Because the VAS task allows for a more continuous rating than the 2AFC task, it is reasonable to assume that listeners may be using both categorical and continuous information simultaneously to form their responses. During VAS judgments, more neural resources might be recruited to allow access to both types of information, resulting in larger ERP amplitudes. While Toscano et al. (2018) demonstrated that gradient cues are represented earlier in the ERP time course (∼N1) than categorical representations (∼P2), Sarrett et al. (2020) suggested gradiency is encoded on a longer time scale, spanning that of the latency window used here. Moreover, the P2 is not solely a response to exogenous acoustics, but an early endogenous response indexing speech discrimination (Alain et al., 2010; Ben-David et al., 2011), auditory object identification (Ross et al., 2013), and category representation (Bidelman et al., 2020; Bidelman et al., 2013). Thus, it is not surprising that P2 changed with task demands, as the response reflects categorical (perceptual) and continuous (acoustic) components. Still, a novel finding is that these early auditory responses beginning at ∼150 ms are influenced by post-perceptual mechanisms that initiate the motor response much later in time (400-800 ms).

As expected, P2 latencies were longer in noisy than clean conditions across both tasks. Noise-related prolongation of the P2 is likely due to decreased neural synchrony due to masking noise (Billings et al., 2009; Kaplan-Neeman et al., 2006; Whiting et al., 1998). However, these latency effects were more prominent in the VAS compared to 2AFC condition. It is unlikely this reflects mere differences in speed of the motor response during VAS labeling since P2 effects were substantially earlier (600-800 ms) than listeners’ RTs. Instead, stronger noise-related changes in VAS may reflect disproportionately augmented sensory effort when performing identification in noise during a graded (VAS) vs. binary (2AFC) task (Bidelman & Walker, 2017; Crowley & Colrain, 2004; Picton & Hillyard, 1974).

More critically, we found that noise-related shifts in ERP latency were behaviorally relevant; smaller latency prolongations were observed for more gradient compared to discrete listeners. This finding supports the notion that making use of continuous cues to decode ambiguous speech is advantageous, as neural timing is less disturbed by noise among more continuous listeners. It is possible that listeners who weigh continuous information more heavily when making perceptual decisions experience less change in stimulus ambiguity. Alternatively, and by the logic above, gradient listeners may also experience reduced attentional load in the presence of noise, accounting for the smaller changes we find in their ERPs.

### 4.4 Gradient listeners have stronger neural activation in _L_STG

Source reconstruction revealed that the P2 effects were attributed to early activation in left-lateralized auditory brain regions. Notably, gradient listeners had stronger neural activations in left STG. Prior work has shown that activation in the _L_STG is greater than that in the _R_STG during speech perception (Ramos Nuñez et al., 2020; Turkeltaub & Branch Coslett, 2010). This left > right asymmetry is largely consistent with theories of brain lateralization (Hickok & Poeppel, 2007) and hemispheric differences in categorization which suggest speech labeling is processed dominantly by the left hemisphere and music labeling by the right (Mankel et al., 2022; Zatorre et al., 1992). Additionally, larger STG responses in graded listeners could reflect increased engagement of working memory resources that help maintain and refresh the neural trace of acoustic-sensory information of the speech signal prior to labeling. Indeed, stronger sustained activity within left (but not right) auditory cortex is observed under more demanding auditory working memory loads as listeners retain verbal sounds in memory (Bidelman et al., 2021; Kumar et al., 2016). Conceivably, the leftward bias in STG source activations we find for gradient listeners could reflect their heavier retention of continuous acoustic information in the auditory sensory-memory buffer prior to assigning a category label.

Non-auditory regions such as inferior frontal gyrus (IFG) have been shown to predict behavioral performance in categorization tasks (Bidelman & Walker, 2019; Golestani & Zatorre, 2004; Lee et al., 2012; Meyers et al., 2008). While we expected to find IFG involvement, the association between behavioral and neural measures was instead restricted to canonical auditory regions (_L_STG). Behavioral tasks such as those used here inherently employ both sensory and decision-making processes. Studies suggest a functional distinction between the two operations whereby activity in auditory cortical regions (including STG) maps onto sound identification (sensory process), while inferior frontal regions map onto reaction time (decision-making process) (Binder et al., 2004; Du et al., 2014). Our findings are consistent with these functional distinctions. Activation of _L_STG was correlated with a measure of precision (i.e., behavioral slope) rather than speed of sound identification. As such, we infer that how a signal is encoded at the level of auditory cortex may predict the degree of categorical perception a listener experiences. In support of this notion, we have shown that neural representations for speech in auditory cortex reorganize to take on more abstract, categorical organization with increased listening experience of the individual (Bidelman & Walker, 2019). Thus, it is possible that continuous feature coding in auditory temporal cortex is initially more effective in supporting speech sound identification (present study) but that over time, intensive auditory or language experience causes it to re-organize (Guenther et al., 2004) and begin supporting abstract phonetic representations for speech in and of itself (Bidelman & Lee, 2015; Bidelman & Walker, 2019; Chang et al., 2010)

## 5. Conclusions

We examined brain and behavioral links between two fundamental operations in speech perception: categorization and speech-in-noise listening skills. ERPs revealed task-dependent changes in early neural responses starting around 150 ms that differentiated categorical from gradient listeners. Gradient listeners had higher SIN comprehension scores, more resilience to noise-related degradation in speech encoding, and stronger neural responses in _L_STG than their discrete/categorical listening peers. While gradient listening was beneficial in multiple domains here, whether our findings extend to more realistic listening environments remains to be investigated. Listeners may adapt their strategies on the fly in real world situations, switching strategies to adapt to changes in signals. Different task structures using more complex stimuli such as sentences or spatially separated streams may find different utility for the gradient strategy. Categorization and SIN deficits are common hallmarks of a variety of auditory-based disorders (Cunningham et al., 2001; Dole et al., 2012; Dole et al., 2014; Lagacé et al., 2010; Putter-Katz et al., 2008; Warrier et al., 2004). Therefore, documenting associations between these skills may provide a linking hypothesis to understand certain communication deficits (Calcus et al., 2016). Future work should examine how a gradient listening strategy could be trained in listeners with poor SIN comprehension to investigate the feasibility of such auditory training as a rehabilitation tool.

## Acknowledgments

This work was supported by the National Institute on Deafness and Other Communication Disorders (R01DC016267 to G.M.B.). Declarations of interest: none.

## Ethical Statement

Participants all provided written informed consent to participate and all procedures were performed in compliance with the Indiana University Institutional Review Board (protocol number 14860, approved April 7, 2022).

## Footnotes

1 This correlation was still significant [*r*(14) = 0.48, *p* = 0.044] when the two participants with negative latency shifts were removed from the analysis.

